# Profiles of Neuroaffective Responding to Natural Reward-Predicting Cues and Alcohol Images in Alcohol Use Disorder: An Event-Related Potential Approach

**DOI:** 10.1101/2025.10.07.680946

**Authors:** Christopher T. Sege, Rhiannon M. Walton, Lisa M. McTeague, Francesco Versace

**Author notes:** <u>DECLARATION OF COMPETING INTERESTS</u>: None to declare.

## Abstract

**BACKGROUND AND AIMS:** Incentive salience attribution (ISA) is the process by which the cues associated with reward take on value themselves. Individual differences in ISA are known to predict reward seeking and consuming behavior in animals and humans, suggesting prognostic potential in use disorders like alcohol use disorder (AUD). Since AUD etiology is thought to involve changes in natural (e.g., food) reward processing as much as alcohol-related processing, current research tested if differences in natural reward cue ISA are apparent, and predictive of behavior, in AUD.

**METHODS:** 30 individuals meeting AUD criteria and reporting heavy alcohol consumption on timeline follow-back (TLFB) participated. Subjects completed a validated ISA task wherein pictures depicting inherently appetitive, aversive, or emotionally neutral content are shown along with pictures that are less evocative but are predictive of reward (candy) delivery and (in this study) also alcohol images. To capture individual variations in ISA, a picture-elicited neural response that scales with picture salience – the late positive potential (LPP) – is indexed and then submitted to *k*-means clustering to classify different modulation profiles.

**RESULTS:** Replicating prior work, clustering indicated a 2-group solution where an “F>P” (n=14) group showed *enhanced* LPPs for food-predicting, relative to standard appetitive, pictures and an “F<P” group (n=16) showed *reduced* LPPs for food-predicting pictures. Also replicating prior work, both groups showed typical enhancement for appetitive and standard aversive, relative to neutral, pictures; and contrary to expectations, both also showed minimal LPP enhancement for alcohol pictures. Examining behavioral correlates of F>P versus F<P status, groups did not differ in reward consumption during the task, nor did they differ in AUD symptoms or weekly drinking per TLFB. Finally, while the LPP was not enhanced for alcohol pictures, enhancement for alcohol relative to neutral pictures was observed for an earlier (P300) neural response – and this was consistent across F>P and F<P groups.

**CONCLUSIONS:** Similarly sized groups of individuals who were more versus less responsive to natural reward cues is consistent with prior studies in non-AUD samples. At the same time, the lack of commensurate differences in task reward consumption differs from those studies, and could reflect change in natural reward processing in individuals with AUD. Implications for continued research are discussed.

## INTRODUCTION

Incentive salience attribution (ISA) is the process whereby cues that predict rewarding outcomes come to have value themselves.^1^ Potentially relevant to clinical assessment, individual differences in ISA to reward-predicting cues have been demonstrated in animals and humans – in animals by measuring cue-related behavior such that, for example, when repeated trials present cues and then deliver rewards in separate areas, an increase in running toward the cue indicates high ISA (“sign tracking”) whereas persistence of cue-induced running to where reward will be delivered indicates low ISA (“goal tracking;” see Flagel et al., 2009^2^). In humans, then, ISA has been tested with electrocortical rather than behavioral measures; and specifically, by testing how a picture-elicited late positive potential (LPP) that tracks picture salience/ survival relevance^3-5^ is or is not also enhanced for otherwise low-salience pictures that predict reward (e.g., candy) delivery.^6-9^ Across studies, ISA differences manifest in cluster analysis-driven identification of one group who shows enhanced LPPs for reward-predicting (compared to inherently salient appetitive, e.g., erotic,^10^ content-depicting) pictures and another who shows *reduced* LPPs for reward-predicting (compared to standard appetitive) images. Indicating behavioral relevance of this finding, these studies also repeatedly find that groups with enhanced reward-predicting image LPPs consume more task rewards than groups without such enhancement – supporting a relationship between a “sign tracking”-like neural response and increased reward consumption in humans as in animals.^7-9^ Building on this even more, studies with clinic samples also relate a sign tracking-like response to obesity severity (in work with obese individuals and food-related images^11^) and treatment outcome (e.g., in work with heavy smokers and smoking-related images^12^). Thus, ISA research together suggests great promise as a prognostic indicator in disorders of problem use.

Despite its promise, ISA has not been tested as reward-consuming behavior or disorder variable predictor in the most prevalent,^13^ public health-burdening^14^ use disorder in the United States – alcohol use disorder (AUD). Informing such investigation, prominent models of AUD posit that pre-morbid general reward-related traits (e.g., reward-seeking, impulsivity) and post-morbid changes in *natural* (e.g., food^15^) reward processing could be as key to AUD etiology ^16,17^ – and perhaps *more* critical to driving relapse risk^18^ – as are disorder-related changes in alcohol reward processing. Inasmuch as establishing behavioral/clinical predictiveness of natural reward cue ISA in AUD could also have the practical implication that clinical ISA assessment might not require alcohol delivery, this study aimed to be the first to test natural reward, rather than disorder-specific, ISA as a use disorder-relevant variable and to do this in a sample of individuals meeting AUD criteria.

In addition to testing natural reward ISA as an AUD-relevant indicator, this study leveraged the fact that ISA tasks use a variety of semantically rich pictures as core stimuli to also test for AUD-related changes in other emotional processing dimensions. In particular, this work was able to simultaneously test for differences in (and predictiveness of) alcohol cue processing by also including disorder-specific alcohol images in our ISA task. In addition, as a standard feature the ISA task also includes normatively appetitive (i.e., pleasant), aversive (i.e., unpleasant), and emotionally neutral images as comparators, and as such can test if disorder-relevant differences are in response to specific (e.g., reward-predictive or alcohol-related) stimuli or instead impact emotional processing more broadly.^19^ Taken as a whole, then, the current study had two critical aims: first, to test for individual differences in, and behavioral predictiveness of, natural reward-related ISA in an AUD sample; and second, to also test for differences in reactivity to alcohol-specific images, and/or general emotion processing, in AUD.

## METHOD

### Participants

Individuals reporting heavy alcohol use (8+ drinks/week for women, 15+ drinks/week for men^20^) and who met DSM-5 AUD criteria were recruited (with an aim toward a similarly sized sample as prior work with clinical samples, i.e. n=∼30^12,21^). Participants were largely recruited and screened by the recruitment core of a National Institute of Alcoholism and Alcohol Abuse (NIAAA) Alcohol Research Center (ARC), while a small number were recruited via broader advertisement across the study institution or word-of-mouth referral. After initial screening by ARC or study staff, diagnosis and consumption were confirmed at the study visit for all subjects.

### AUD Characterization

#### Diagnostic Confirmation – MINI Interview

The Mini International Neuropsychiatric Interview (M.I.N.I.) for DSM-5^22^ was conducted to confirm AUD and rule out exclusions (manic or psychotic episode history). All participants endorsed diagnostic criteria for AUD of at least mild severity. Moreover, no participant met criteria for a lifetime manic or psychotic episode.

#### Dimensional AUD Symptoms – Alcohol Use Disorder Identification Test (AUDIT)

The AUDIT^23^ is the most used questionnaire to measure dimensional AUD symptoms. The AUDIT comprises 10 questions quantifying how often people use alcohol problematically and experience related symptoms (e.g., craving, difficulty stopping). Items 1 through 8 are rated from 0 (never) – 4 (4+ times/week), while item 9 (“have you or someone else been injured as a result of drinking”) and 10 (“has a relative, friend, doctor or other health worker been concerned about/advised you to cut down your drinking”) are rated as 0 (no), 2 (yes, but not in the last month), or 4 (yes, in the last month). Along with its common use in research, the AUDIT is widely used in clinic settings to screen for alcohol use disorder – and as such, cut-off scores for low-risk consumption (1–7), hazardous/harmful alcohol use (8–14), or likely alcohol dependence (15+) have been established by the World Health Organization.^24^

#### Alcohol Use – Timeline Follow Back (TLFB)

The TLFB^25^ is a commonly used standardized interview methodology for quantifying alcohol consumption over a set period (in this study, the past 30 days). In TLFB procedures, participants are first oriented to standard drink amounts (12-ounce beer at 5% alcohol by volume [ABV]; 5-ounce wine/beer at 12% ABV; 1.5-ounce liquor shot at 40% ABV). Next, participants are shown the past 30 days on a calendar, oriented to weekends/holidays during that span, and asked to quantify number of drinks consumed each day starting from the day before the visit and working backward to 30 days prior. TLFB procedures produce a reliable estimate of typical current alcohol consumption for research and clinical purposes; and by recalculating to produce a weekly average, it can determine if a person falls above NIAAA heavy consumption definitions (8+ drinks/week for women, 15+ drinks/week for men^20^).

### Incentive Salience Attribution (ISA) Task

As a primary assay of emotional and alcohol cue processing, participants completed a modified ISA task developed by Versace et al.^6,7^ In this task, participants passively view pictures that vary in the motive salience (appetitive, aversive, emotionally neutral) and intensity (high arousal, low arousal) of content depicted. Along with inherently appetitive and aversive images, participants also see a typically less evocative content category (sweet foods, e.g. candies, cakes^26^) that, after 2 seconds, are superimposed upon by a word (“M&M”) indicating the person can retrieve an actual food reward (M&M candy) from a bowl on their right and chose to eat or discard it. After eating the M&M or discarding it into a bowl on the left, participants push a button indicating their choice and this removes the food image (after minimum 4-second presentation) and continues the task. On all other trials, pictures are not followed by food reward but, instead, simply presented for 4 seconds and then removed from the screen. After all trials, a 0.5–2.5-second inter-trial interval precedes the next trial.

In this study, participants sat 100cm from a computer monitor and viewed 16.875cm x 22.5cm (4.8° x 6.4° visual angle from center) pictures. Most pictures were the images from the International Affective Picture System (IAPS^27^) that were also used in initial work^7^ to depict: 1) sweet foods; 2) erotica; 3) romantic scenes; 4) neutral people (e.g., everyday conversations); 5) neutral objects (e.g., stapler); 6) aversive objects (e.g., toxic waste barrels); 7) violent scenes, and; 8) mutilated bodies/bodily fluids. As described in initial work,^7^ normative ratings from the IAPS^27^ indicate that erotic and romantic images represent high- and low-arousal appetitive (pleasant) contents respectively, mutilation and violent images represent high- and low-arousal aversive (unpleasant) contents respectively, and food and aversive objects are like neutral scenes in salience and intensity.

Along with images from initial work, in this study alcohol images from the Wake Forest Alcohol Imagery Set (WFAIS^28^) were also included. Normative WFAIS^28^ data suggest people without problem alcohol use rate alcohol images as neutral and low intensity, while individuals with problem use rate these images as more pleasant and arousing. Here, alcohol images were intermixed throughout the task and all contents were presented in random order except that no more than two from the same category could be presented consecutively. 330 total pictures (60 reward-predicting and alcohol pictures, 30 of other contents) were shown over the entire task.

### Electroencephalography

#### Online Data Collection

To index event-related potential indicators of image processing, electroencephalography (EEG) was recorded with a Brain Products© actiCHamp® 32-channel active-channel system. ActiCap© cloth caps held sensors in 10–20 standard positions on the scalp. Data were sampled at 500Hz and referenced to a common mode and then sensor Fz. Impedances were kept below 15 kOhms and data were not filtered online except for a 250-Hz anti-aliasing filter. Along with EEG, electrooculogram (EOG) data (also sampled at 500Hz) were collected with Ag/ AgCl sensors above and below the right eye and laterally to each eye.

#### Offline (Post-Task) Data Processing

After collection, processing with Brain Vision Analyzer® replicated parameters in the prior studies as much as possible^6-9^. The exception was re-referencing, which in this study was to linked mastoids (TP9, TP10) rather than average reference due to less sensor density than in prior work. After re-referencing, EEG was filtered with Butterworth 1/3-amplitude 0.1Hz (Order 4) – 30Hz (Order 8) cut-off filters and segmented from 100ms pre-onset – 1000ms post-onset of each picture. Next, eye movement artifacts were removed via independent components analysis decomposition of combined EEG/EOG data,^29^ and other artifacts were then removed with extreme voltage (>=|100|µV), extreme voltage shift (>=100µV within the segment), extreme voltage step (>=25µV from sample to sample), and voltage non-variation (<=.50 µV variation for >100ms) criteria.^6-9,30^ Artifact correction was used to identify bad channels that were then interpolated using spherical splines^31^ unless >2 bad channels occurred in close proximity or >20% of channels were bad on a trial (in which case the trial was removed^32^). Artifact correction led to 4 (13.3%) trials/condition removed on average, with no difference in trials removed across contents, *F*(4,116)=1.5, *p*=.21, η_*p*_ ^*2*^=.05). After correction, each EEG segment was baseline (100ms-0ms) corrected and averaged across trials within each picture content.

Processing steps described above were used to extract characteristic components of the picture-evoked event-related potential, including a prominent positive slow-wave component – the late positive potential (LPP) – that tracks image salience.^3-5,26^ As in prior work, the LPP was quantified for analysis here by averaging EEG in the observed time/sensor window of maximal LPP amplitude across contents; with prior studies, including those with the ISA task,^6-9,11,12^ indicating canonical LPP voltage maxima in a ∼300ms to ∼1000ms window after picture onset and at central/parietal EEG sensors (see also Bradley et al.^26^). For comparison to the canonical observation, time/sensor windows observed for the LPP in this study are presented in Results.

While prior studies of ISA and of broader emotional picture processing have focused on the LPP as a main event-related potential component of interest, some studies of alcohol image processing^33^ indicate differences for a related earlier P300 component that is commonly thought of as a marker of initial salience detection.^34^ Given a possibility that alcohol effects could be specific to this component, P300 was also extracted for analysis by averaging within the observed time/sensor window for this component; with canonical observation of maxima for this component in a ∼250ms – 450ms window after picture onset and broadly across central sites. As for the LPP, actual observed time/sensor windows for the P300 are presented in Results.

### Procedure

After consent, participants completed interviews (M.I.N.I., TLFB) and the AUDIT, and they rated their pre-task hunger on a visual analog scale from “Greatest imaginable hunger” (value 0) to “Greatest imaginable fullness” (value 40; value 20 = “Neither hungry nor full”). Next, EEG sensors were placed and task instructions given. Before the main task, participants completed a short practice block (with instructions to not take or eat candies) to orient them. Completing the full ISA task took ∼35 minutes on average.

After the task, participants rated hunger again and then pleasantness of picture contents and M&M rewards. For pictures, participants rated pictures of “food,” “erotic couples,” “clothed couples,” “alcohol,” “everyday objects (e.g., a rolling pin),” “everyday scenes (e.g., two people talking),” “pollution,” “violence,” and “mutilated bodies or bodily fluids” on a Likert-type 1 (*very unpleasant*) – 9 (*very pleasant*; 5 = *neither pleasant nor unpleasant*) scale. For M&Ms, subjects rated how much they “like or want/wanted M&M candies” both “usually” and “during the task” (separate ratings) on a 1 (*not at all*) – 9 (*very much so*) scale. Finally, sensors were removed and participants debriefed and paid.

### Data Analysis

Primary analyses examined general LPP modulations by picture salience and then individual differences in this modulation and how they related to reward-consuming behavior and AUD/comorbid symptomatology and drinking. In the main article, task-based analyses focused on superordinate standard appetitive (erotica and romance average), neutral (people and objects average), standard aversive (violence and mutilation average), food-predicting, and alcohol categories. For clarity, further analysis of picture sub-contents are then presented in supporting information. In all analyses, picture category/content is a within-subjects factor.^35.36^ Using this approach, then, analyses first tested task effects for LPP amplitudes and also picture ratings with Analyses of Variance (ANOVAs) and indicated follow-up *t*-tests.

Following confirmation of general task effects, individual differences in natural reward-related ISA were examined in an problem alcohol-using sample by replicating neural response classification methods from prior work.^7-9,11,12^ Specifically, LPP modulation profiles were grouped using *k*-means clustering as implemented in a JmP© statistical package. In *k*-means clustering, select variables are input to a multivariate algorithm that generates a user-specified number of groups (clusters) that minimize in-cluster variability and maximize between-cluster variability.^37^ Since the *k*-means algorithm is unsupervised, clustering in this manner is data-driven with the only constraints being selection of input data and number of user-specified groups. Regarding the latter, examination of a cubic clustering criterion (CCC) in JmP^38^ methods can then be used to determine optimal cluster number. To replicate and extend testing of natural reward-related ISA variations to an AUD sample, *k*-means clustering with CCC evaluation was done on appetitive, neutral, aversive, and food-predicting LPPs (*z*-score normalized within-subject to remove overall level differences^39^), with alcohol LPPs not included so that methods replicated prior work. After clustering, derived groups were compared for LPP modulations to characterize effects, and for M&M reward consumption/ratings, picture pleasantness ratings, alcohol-related variables (TLFB, AUDIT), and comorbid (anxiety/depression) symptom variables to determine how clustering predicted reward consumption and clinical variables in this sample.

Finally, in addition to replication of LPP analyses to characterize emotional picture processing and individual differences therein, modulation of an earlier P300 event-related potential component was also tested in this problem alcohol-using sample. T2o explore potential individual differences and predictiveness of this component, P300 differences were compared across groups derived via *k*-means clustering and P300 amplitude scores were also correlated with behavioral and clinical variables.

## RESULTS

### Sample

The final sample was 30 individuals with mean age 43.0 years (SD=16.5), an equal number of biological men and women, and predominantly (n=25, 83.3%) non-Hispanic White race/ethnicity. Mean AUDIT score fell within a WHO-defined hazardous range for women (M=13.7; SD=5.6) and men (M=14.7; SD=5.6), and TLFB-assessed drinks/week were above NIAAA heavy drinking cut-offs for women (M=20.0, SD=11.3) and men (M=30.7, SD=13.8).^*1*^

### Task Behavior and Ratings

Across the sample, participants ate an average of 13 (SD=14.1) M&Ms over the course of the task. Participants generally rated moderate hunger (M=20.2; SD=5.4) prior to the task, and this did not change after the task (M=21.0; SD=6.2), *t*(27)=-0.8, *p*=0.5, *d*=-0.1. After the task, participants rated moderate general liking of M&Ms (M=5.4; SD=2.3) and wanting of M&Ms during the task (M=4.1; SD=2.2).

Regarding task picture ratings, overall modulation patterns, *F*(4,116)=125.6, *p*<.001, *η*_*p*_^*2*^*=*.81, were consistent with prior research for standard images such that appetitive images were rated as more pleasant than neutral, *t*(29)=5.1, *p*<.001, *d*=.93, and aversive images as more unpleasant than neutral *t*(29)=-18.2, *p*< .001, *d*=-3.3 (see **Figure 1**). Next, both reward-predicting, *t*(29)=5.4, *p*<.001, *d*=1.0, and alcohol-related, *t*(29)=3.8, *p*<.001, *d*=.69, pictures were also rated as more pleasant than neutral, and neither reward-predicting, *t*(29)=1.6, *p*=.12, *d*=.29, nor alcohol, *t*(29)=-1.3, *p*=.22, *d*=-.23, differed from standard appetitive images. Finally, reward-predicting images were more also rated as more pleasant than alcohol, *t*(29)=2.8, *p*=.01, *d*=.50.

**Figure 1.**
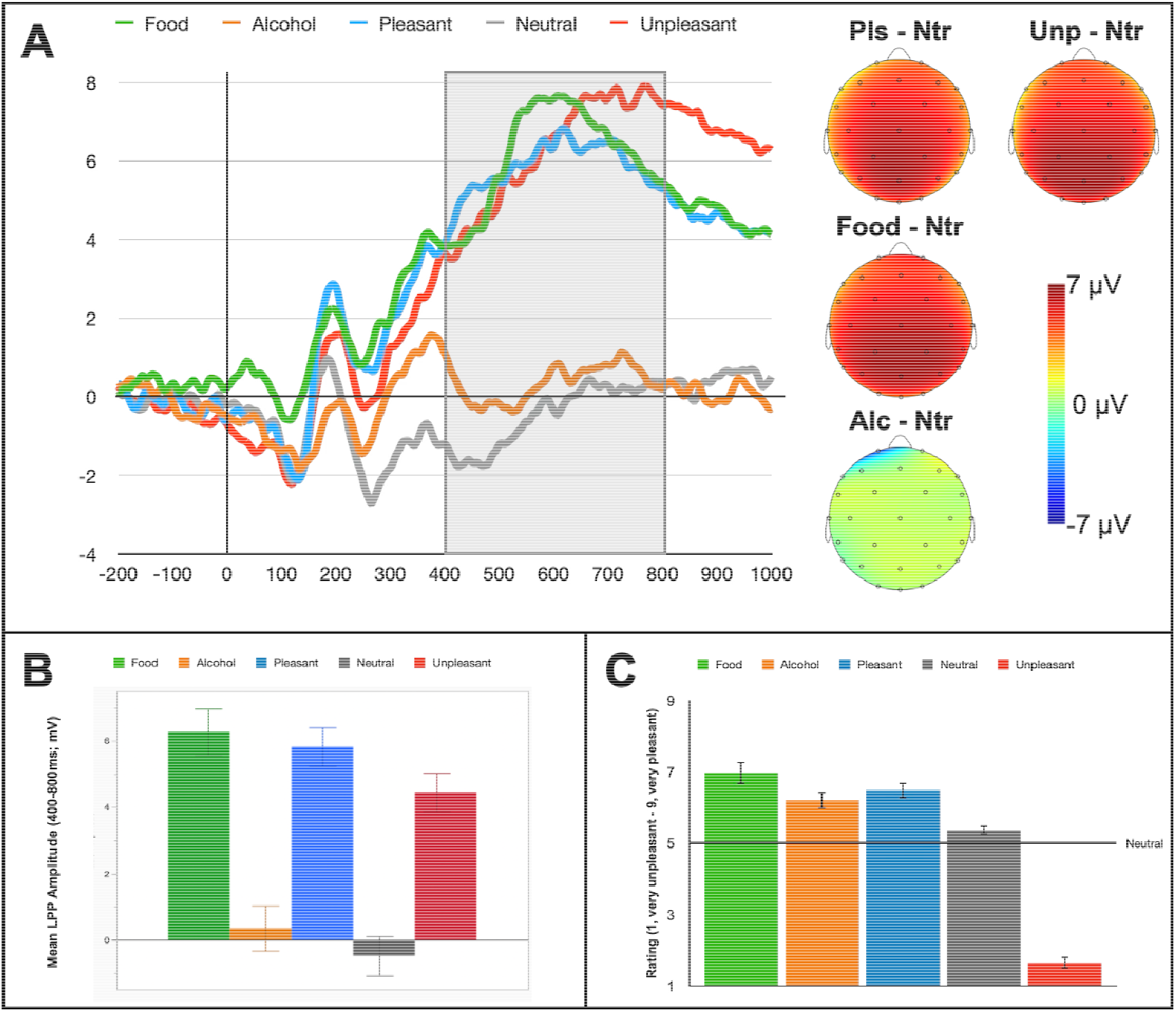
Panel A, Event-related potential to images in the Incentive Salience Attribution task. In the waveform plot, 0 indicates picture onset and shaded area highlights the window of peak late positive potential (LPP) component amplitude. Scalp topographies show the mean of LPP amplitude differences from neutral for each other image type within the 400 – 800ms window. Panel B, bar chart representation of LPP amplitude averaged within a 400 – 800ms time window and across centroparietal (Cz, CP1, CP2, Pz) sites for each picture category. Panel C, average rated pleasantness for each picture category.

### Late Positive Potential (LPP) Analyses

#### Task Effects

Across the sample, LPP morphology was in line with prior observations with amplitudes maximal from 450–750ms and at centroparietal sites (Cz, CP1, CP2, Pz, P3, P4; **Figure 1**). Robust modulation across contents, *F*(4,116)=50.7, *p*<.001, *η*_*p*_^*2*^*=*.64, was apparent such that, first, standard appetitive, *t*(29)= 9.5, *p*<.001, *d*=1.7, and aversive, *t*(29)=9.7, *p*<.001, *d*=1.8, image LPPs were each enhanced compared to LPPs for neutral images, and appetitive was also enhanced relative to aversive, *t*(29)=3.5, *p*=.002, *d*=0.6, LPPs. Next, reward-predicting image LPPs were also larger than neutral, *t*(29)=9.7, *p*<.001, *d*=.1.8, and aversive, *t*(29)=3.3, *p*=.003, *d*=0.6, LPPs, and they did not differ from appetitive LPPs, *t*(29)=0.7, *p*=.47, *d*=.13.

Finally, alcohol-related LPPs did not significantly differ from neutral, *t*(29)=1.8, *p*=.08, *d*=.33, and they were smaller than LPPs for appetitive, *t*(29)=-7.5, *p*<.001, *d*=-1.4, aversive, *t*(29)=-7.3, *p*<.001, *d*=-1.3, and reward-predicting, *t*(29)=-7.6, *p*<.001, *d*=-1.4, images.

#### Individual Difference Classification

*K*-means clustering of LPP profiles indicated a two-cluster solution that captured the data. Modulation differences across groups, Group X Valence *F*(4,112)=9.0, *p*<.001, *η*_*p*_^*2*^*=*.24, were specifically reliable for reward-predicting images such that: 1) an “F>P” group (n=14) had larger LPPs for reward-predicting images than for neutral, *t*(13)=9.8, *p*<.001, *d*=2.6, appetitive, *t*(13)=6.4, *p*<.001, *d*=1.7, and aversive, *t*(13)=5.6, *p*<.001, *d*=1.5, images, and; 2) an “F<P” group (n=16) still had larger reward-predicting as compared to neutral LPPs, *t*(15)= 7.0, *p*<.001, *d*=1.8, but also had *smaller* reward-predicting than appetitive LPPs, *t*(15)=-3.6, *p*=.003, *d*=-0.9, and no difference between reward and aversive LPPs, *t*(15)=-0.2, *p*=.81, *d*=-0.1 (**Figure 2**). Across other categories, modulation patterns were similar for each group; and as such, when groups were directly compared differences arose for reward-predicting images, *t*(28)= 3.7, *p*<.001, *d*=1.4, but not for appetitive, *t*(28)=-2.0, *p*=.06, *d*=-.72, neutral, *t*(28)=-0.5, *p*=.61, *d*=0.2, or aversive, *t*(28)=-0.4, *p*=.36, *d*=-.13, LPPs. Finally, when alcohol pictures were tested, groups did not differ from each other, *t*(28)=-0.3, *p*=.80, *d*=-0.1, and alcohol did not differ from neutral for F>P, *t*(14)=1.6, *p*=.12, *d*=.44, or F<P, *t*(14)=1.0, *p*=.33, *d*=.25, groups.

**Figure 2.**
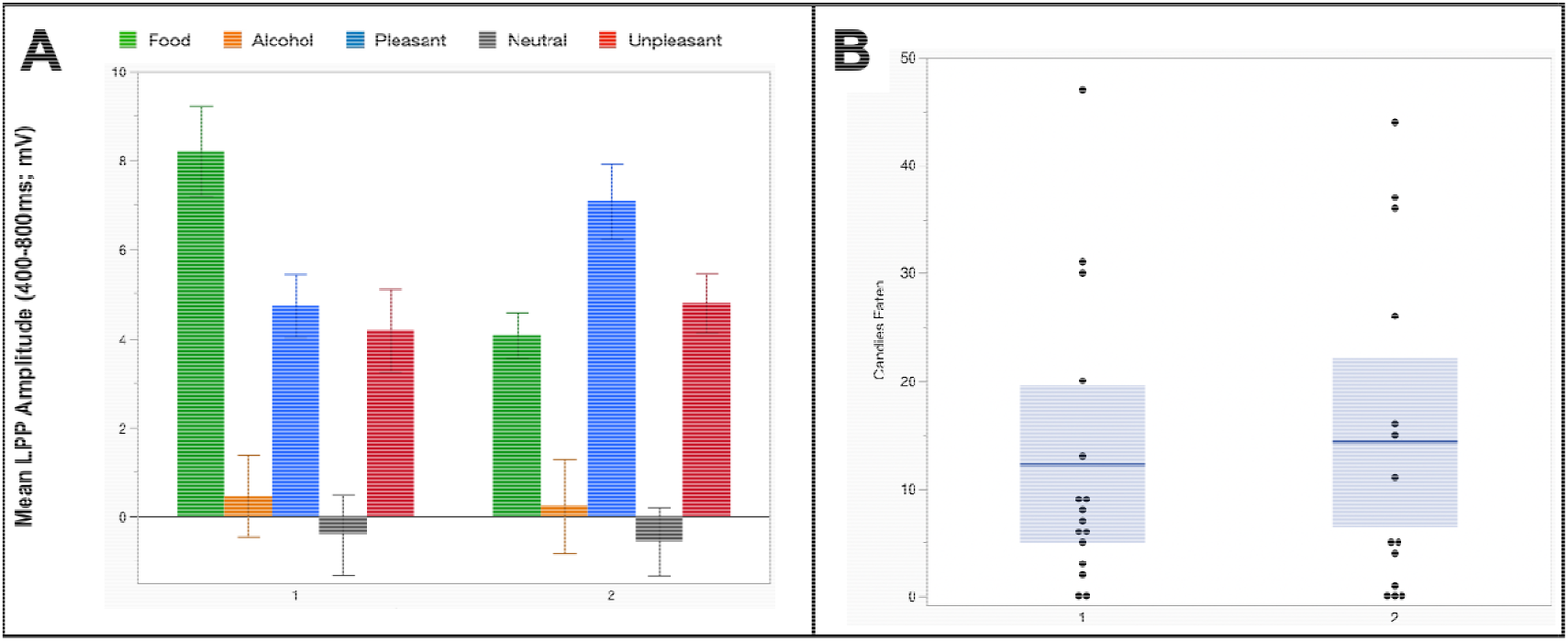
Panel A, mean LPP amplitudes for each picture category across *k*-means clusterderived “Food>Pleasant (F>P)” and “Food<Pleasant (F<P)” groups. Panel b, number of candies eaten by each participant across *k*-means-derived groups. In each column, the blue line and shaded area represent mean and variance for that group and each dot represents a participant.

Next, testing how LPP-based grouping predicted task behavior (**Figure 2** shows candy consumption and **Table 1** shows M&M, hunger, and picture ratings) revealed no differences for any variable; such that, first, F>P and F<P groups did not differ in the number of candies eaten during the task, *t*(28)=-0.6, *p*=.53, *d*=-.23, wanting of M&Ms during the task, *t*(28)=-0.4, *p*=.69, *d*=-.15, or general liking of M&Ms, *t*(28)=0.9, *p*=.39, *d*=.32. Next, there were also no differences in hunger before, *t*(28)=0.5, *p*=.63, *d*=.19, or after, *t*(28)=-0.3, *p*=.78, *d*=-.11, the ISA task.

**Table 1.**
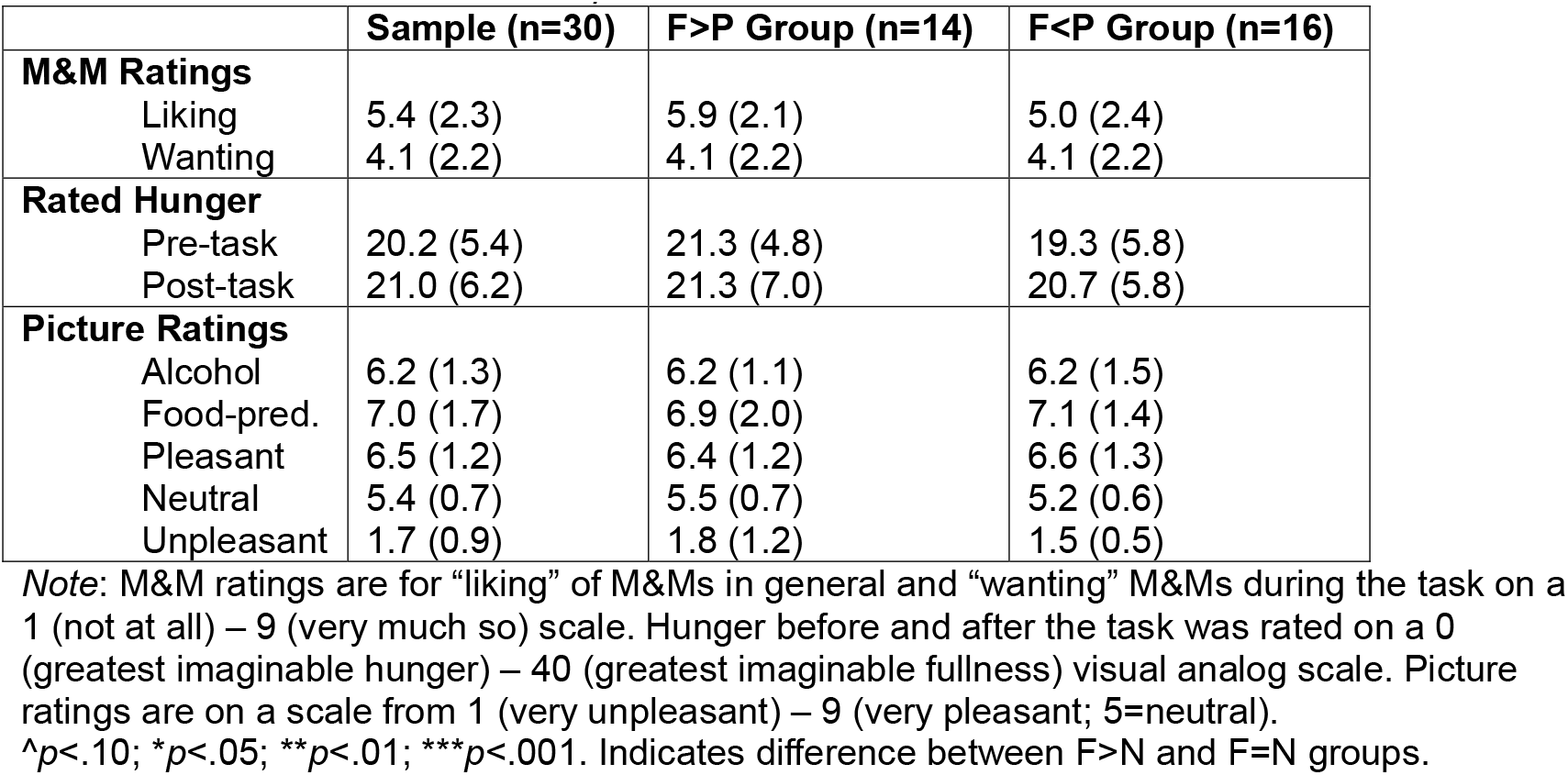
Means (Standard Deviations) of Task-related Behavioral Variables Across K-Means Cluster-Derived F>P and F<P Groups.

|  | Sample (n=30) | F>P Group (n=14) | F<P Group (n=16) |
| --- | --- | --- | --- |
| **M&M Ratings** |  |  |  |
| Liking | 5.4 (2.3) | 5.9 (2.1) | 5.0 (2.4) |
| Wanting | 4.1 (2.2) | 4.1 (2.2) | 4.1 (2.2) |
| **Rated Hunger** |  |  |  |
| Pre-task | 20.2 (5.4) | 21.3 (4.8) | 19.3 (5.8) |
| Post-task | 21.0 (6.2) | 21.3 (7.0) | 20.7 (5.8) |
| **Picture Ratings** |  |  |  |
| Alcohol | 6.2 (1.3) | 6.2 (1.1) | 6.2 (1.5) |
| Food-pred. | 7.0 (1.7) | 6.9 (2.0) | 7.1 (1.4) |
| Pleasant | 6.5 (1.2) | 6.4 (1.2) | 6.6 (1.3) |
| Neutral | 5.4 (0.7) | 5.5 (0.7) | 5.2 (0.6) |
| Unpleasant | 1.7 (0.9) | 1.8 (1.2) | 1.5 (0.5) |

*Note:* M&M ratings are for “liking” of M&Ms in general and “wanting” M&Ms during the task on a 1 (not at all) – 9 (very much so) scale. Hunger before and after the task was rated on a 0 (greatest imaginable hunger) – 40 (greatest imaginable fullness) visual analog scale. Picture ratings are on a scale from 1 (very unpleasant) – 9 (very pleasant; 5=neutral).
<sup>^</sup> $p < .10$ ; \* $p < .05$ ; \*\* $p < .01$ ; \*\*\* $p < .001$ . Indicates difference between F>N and F=N groups.

Finally, analysis of picture ratings did not indicate an interaction, *F*(4,112)=1.7, *p*=.16, *η*_*p*_^*2*^*=*.06, nor did group differences arise in the pleasantness of reward, *t*(28)=-1.0, *p*=.34, *d*=-.36, alcohol, *t*(28)=-0.2, *p*=.82, *d*=-.08, appetitive, *t*(28)=-1.6, *p*=.12, *d*=-.59, aversive, *t*(28)=1.0, *p*=.33, *d*=.36, or neutral, *t*(28)=1.3, *p*=.21, *d*=.47, images.

Next, when ISA groups were compared for clinical variables (**see Table 2**), AUDIT, *t*(28)= -0.1, *p*=.91, *d*=-.04, and TLFB weekly drinking, *t*(28)=0.8, *p*=.44, *d*=.29, were similar across groups. Finally, comparing sample characteristics indicated no group differences in biological sex, _χ_^2^(29)=0.5, *p*=.46, race/ ethnicity, _χ_^2^(29)=0.7, *p*=.39, or age, *t*(28)=-0.8, *p*=.42, *d* = -.31.

**Table 2.** Sample Characteristics, Including Means (Standard Deviations) of AUD-Related Variables, across K-Means Cluster-Derived F>P and F<P Groups.

|  | Sample (n=30) | F>P Group (n=14) | F<P Group (n=16) |
| --- | --- | --- | --- |
| **Sex (N[%] Male)** | 15 (50.0) | 8 (57.1) | 7 (43.8) |
| **Race/ Ethnicity (N[%] White)** | 25 (83.3) | 12 (85.7) | 13 (81.3) |
| **Age** | 43.0 (16.5) | 44.5 (17.5) | 41.7 (16.1) |
| **AUDIT Total Score** | 14.2 (5.5) | 14.3 (4.9) | 14.2 (6.2) |
| **TLFB Average** | 25.4 (13.5) | 27.8 (16.4) | 23.3 (10.5) |

*Note:* TLFB Average presents average number of alcoholic drinks over 1 week (7 days).
\*Mann-Whitney *U* test = 153.5, $p = .04$ ; Mann-Whitney presented due to deviations from normality in both groups.

### P300 Analyses

#### Task Effects

Visual examination of the morphology of an earlier P300 picture processing ERP component showed a distinct peak maximal from 280–420ms and across central sites (FC1, FC2, Cz, CP1, CP2; **Figure 3**) for neutral, reward-predicting, and alcohol-related images. For standard appetitive and aversive images, amplitudes in this window were enhanced but not clearly distinct from the LPP. Examining P300 amplitudes across contents revealed modulation that was similar to the LPP for standard images, *F*(4,116)=27.7, *p*<.001, *η*_*p*_^*2*^*=*.49 – such that P300s for appetitive, *t*(29)= 6.7, *p*<.001, *d*=1.2, and aversive, *t*(29)=5.4, *p*<.001, *d*=1.0, images were each enhanced compared to neutral, and appetitive was also enhanced compared to aversive, *t*(29)=5.8, *p*<.001, *d*=1.2. Also similarly to LPP, for reward-predicting pictures the P300 was larger than neutral, *t*(29)=8.2, *p*<.001, *d*=.1.5, and aversive, *t*(29)=5.2, *p*<.001, *d*=0.9, but it did not differ from the appetitive P300, *t*(29)=0.5, *p*=.62, *d*=0.1. Finally, unlike the LPP, alcohol P300s were also enhanced relative to neutral, *t*(29)=5.5, *p*<.001, *d*=0.9, and did not differ from aversive, *t*(29)=-0.3, *p*=.79, *d*=-.05, though they were still smaller than appetitive, *t*(29)=-4.0, *p*<.001, *d*=-0.7, and reward, *t*(29)=-4.8, *p*<.001, *d*=-0.9, P300s.

**Figure 3.**
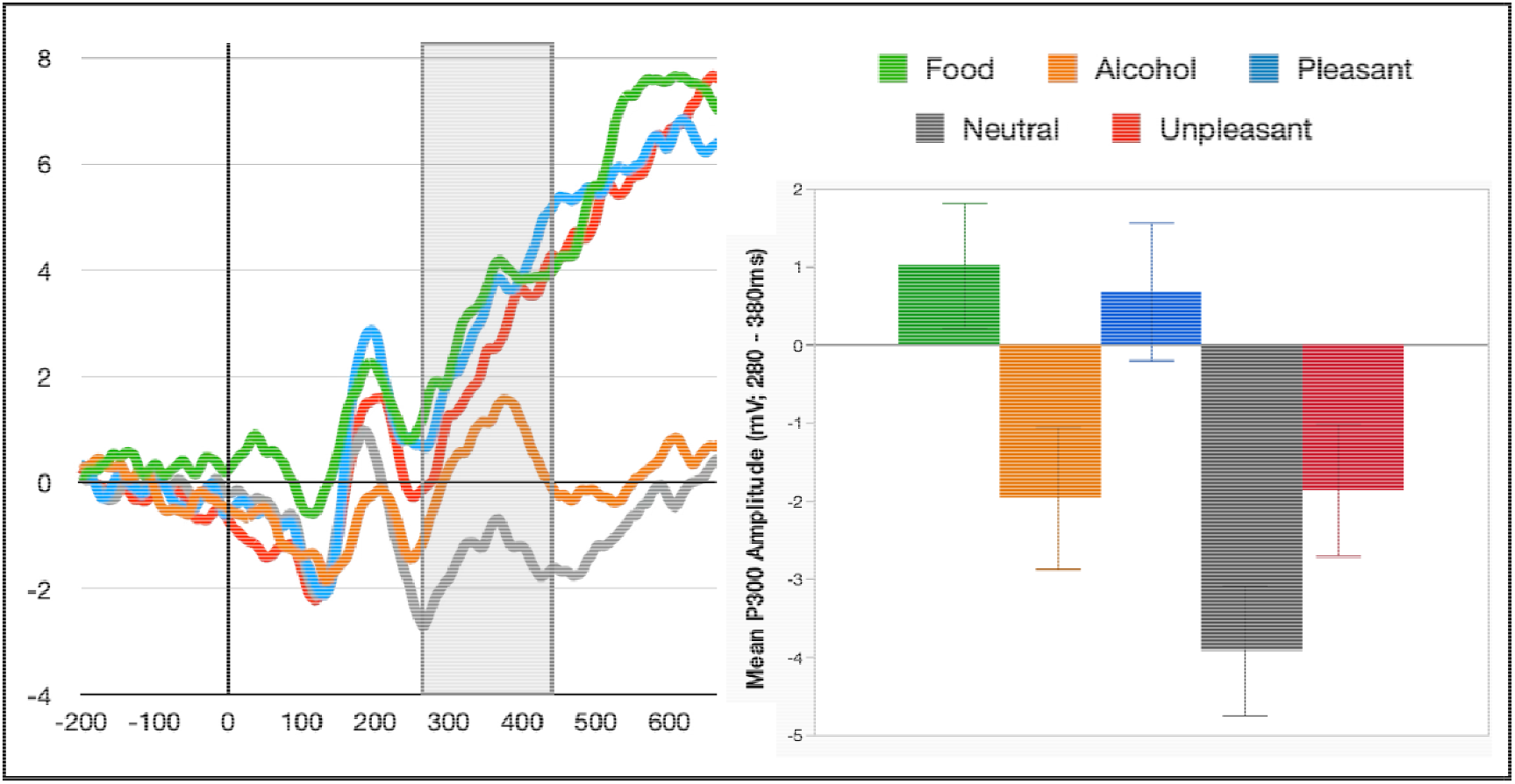
Event-related potential with the window of peak P300 amplitude highlighted. Bar chart represents P300 amplitude averaged within a 280 – 380ms time window and across fronto-central (FC1, FC2, Cz, CP1, CP2) sites for each picture category.

#### Individual Difference Exploration

When P300 modulations were compared across groups derived via *k*-means clustering of the LPP, groups did not differ in P300-interval amplitudes for any content (see **Table 3**). In addition, the F>P and F<P group each showed P300 enhancement compared to neutral for alcohol-related (F>P *t*(13)=4.2, *p*=.001, *d*=1.1; F<P *t*(15)=3.6, *p*=.003, *d*=0.9), reward-predicting (F>P *t*(13)=7.3, *p*<.001, *d*=1.9; F<P *t*(15)=4.9, *p*<.001, *d*=1.2), appetitive (F>P *t*(13)=5.1, *p*<.001, *d*=1.4; F<P *t*(15)=4.7, *p*<.001, *d*=1.2), and unpleasant (F>P *t*(13)=5.6, *p*<.001, *d*=1.5; F<P *t*(15)=3.0, *p*=.008, *d*=0.8), pictures.

**Table 3.** P300 amplitude modulation across K-Means Cluster-Derived F>P and F<P Groups.

|  | Sample (n=30) | F>P Group (n=14) | F<P Group (n=16) | Group Comparison |
| --- | --- | --- | --- | --- |
| | M (SD) | M (SD) | M (SD) | $p$ ($d$) |
| **Picture Content** |  |  |  |  |
| Reward | 1.0 (.80) | 1.5 (1.4) | 0.6 (.91) | .57 (-.21) |
| Alcohol | -2.0 (.91) | -2.2 (1.5) | -1.7 (1.1) | .79 (.10) |
| Pleasant | 0.7 (.88) | -0.5 (1.2) | 1.7 (1.3) | .22 (.46) |
| Neutral | -3.9 (.83) | -4.4 (1.5) | -3.5 (.94) | .58 (.21) |
| Unpleasant | -1.9 (.85) | -2.3 (1.5) | -1.5 (.96) | .66 (.16) |

Next, to further explore potential relationships between alcohol use and P300 given that this component, unlike LPP, was enhanced for alcohol images, potential differences between lower (TLFB weekly average ≦ 20; n=14) vs. higher (TLFB > 20; n=16) drinking groups. In this analysis, drinking groups did not differ in response to alcohol-related pictures, *t*(28)=1.6, *p*=.11, *d*=0.6, or to food-predicting, *t*(28)=0.5, *p*=.60, *d*=0.2, appetitive, *t*(28)=1.1, *p*=.29, *d*=0.4, aversive, *t*(28)=1.6, *p*=.12, *d*=0.6, or neutral, *t*(28)=1.5, *p*=.14, *d*=0.6, pictures. In addition, enhanced P300 for alcohol, relative to neutral, pictures was apparent for the lower-drinking, *t*(14)=3.6, *p*=.003, *d*=1.0, and higher-drinking, *t*(14)=4.0, *p*=.001, *d*=1.0, group alike.

## DISCUSSION

As its first aim, this study tested natural reward cue ISA individual differences – and their behavioral predictiveness – in alcohol use disorder. As a first key finding, cue-related ISA variations were akin to those from other samples – such that half the sample showed enhanced LPPs for reward-predicting cues relative to standard appetitive pictures and half did not. This pattern of individual differences in response to reward-predicting cues is akin to those described in non-human animals – where the propensity to demonstrate increasing responding to the cues that predict reward, rather than just to rewards themselves, has been called “sign tracking” behavior.^2,11^ Similarity of natural reward-related ISA difference patterns in AUD as in other samples could then suggest that this is not a pre-morbid AUD risk predictor – inasmuch as, if it were, disproportionate sign tracker representation might be expected (*cf*. Flagel et al., 2009^2^).

As a second and surprising finding, in the current AUD sample individuals with enhanced LPP to reward-predicting cues did not differ from individuals without this enhancement in the number of candies eaten during the task – an unexpected result given consistent differences that have been observed repeatedly in prior work where enhanced reward cue LPP is associated with increased reward consumption.^6-9^ Given evidence that AUD can alter taste perception and preferences especially of sugary foods,^39^ one interpretation of this finding could be that AUD-induced change in response to natural candy rewards altered the relationship between cue-related ISA and candy consumption. At the same time, with the second aim of this study being to test natural reward cue ISA as a predictor of AUD-specific outcomes, it was also observed that reward cue LPP-based grouping did not predict differences in AUD symptomatology or TLFB-assessed alcohol consumption. Taken as a whole then, it could be that natural reward cue ISA is less predictive of reward consuming behavior in general than would be *alcohol* reward cue ISA – a possibility that must be tested in future research where cues predict alcohol, rather than natural food, rewards.

Finally, in addition to examining individual differences in reward cue ISA this study also tested for potential individual differences in response to alcohol pictures, and/or in general emotional processing, as predictors of behavior in an AUD sample. Regarding alcohol image responsivity, we first observed that LPP response to these images was quite minimal overall – a finding which could be due to alcohol images becoming less salient when presented along with images that predicted actual delivery of the depicted reward. At the same time, enhanced responding to alcohol images was still observed in an earlier salience-sensitive ERP component – the P300 – and this was consistent across reward cue processing groups and exploratory-examined lower vs. higher drinking groups to suggest overall reliability of enhanced alcohol-related P300s in this overall heavy drinking sample. Regarding general processing of standard emotional contents, then, typical enhancements of both P300 and LPP for both appetitive and aversive, compared to neutral, images were consistently observed across the individuals with AUD recruited here. As a whole, then, current results suggest a consistent enhancement of early neural responding to alcohol images in heavy drinking individuals, and also consistently intact emotional processing regardless of whether the response to reward-predicting cues was enhanced or not.

In all, current results suggest that reward-predicting cue ISA has the most promise as a potential individual difference metric for use in clinical settings. Before this can be implemented, though, continued research must address limitations of current work that leave still-open questions. Most critically, future work should compare natural reward-related ISA with ISA to cues that signal *disorder-relevant* (i.e., alcohol) rewards as predictors of relevant behaviors including task reward consumption – an especially important goal given findings in nicotine use disorder that cue-related ISA does still positively predict task reward consumption when the reward is disorder-relevant (vaping).^9^ Additionally, future work should compare results with other non-alcohol rewards (e.g., savory foods, money) to confirm if current findings do represent a change in non-alcohol reward processing generally. Finally, future work should relate ISA (natural and disorder-relevant) with other AUD prognostic variables (e.g., treatment response, relapse risk) to confirm utility of ISA as an outcome predictor that can inform treatment-enhancing individualization strategies.

Even before future work, though, current research contributes to understanding natural reward-related ISA as an AUD-relevant indicator by suggesting 1) similar ISA differences, but also a potentially reversed relationship with task behavior, in an AUD sample; and 2) that ISA and alcohol image responsivity may predict distinct AUD dimensions. Replicating/extending these results could have great impact by supporting development of objective assessments that inform treatment planning – and such assessment could be especially useful if natural reward-related ISA continues to be predictive, obviating necessity to administer the problem stimulus.

## ACKNOWLEDGEMENTS

We thank study participants for contributing their time for this study. Along with Rhiannon Walton who coordinated a majority of the study sessions, we thank study team members Samantha LaPorta and Christina Marsicano for their additional assistance with recruitment and data collection. We thank the Medical University of South Carolina’s Alcohol Research Center – and especially Drs. Howard Becker, Patrick Mulholland, Jim Prisciandaro, and William Mellick – for supporting this research.

## Footnotes

1 women who initially screened positive had TLFB-assessed average drinks/week=6 at time of participation. Excluding these women did not alter statistical patterns in any instance. No man fell below the TLFB cut-off at time of participation (minimum score = 15/week, 1 participant).

